# Population-level learning of non-additive effects enables accurate and efficient prediction of combinatorial perturbation responses

**DOI:** 10.64898/2026.11.20.768136

**Authors:** Shaokun An, Qiyu Gong, Kai Cao

## Abstract

Combinatorial perturbations are widely used to study genetic interactions and to develop drug combinations, but the number of possible combinations grows quadratically with the number of perturbations, and only a small fraction can be measured in a screen experiment. Despite their growing sophistication, deep-learning and foundation models often fail to outperform a simple additive baseline based on measured single-perturbation responses. This suggests a simpler approach that builds on these measurements and directly learns deviations from additivity at the population level. We therefore developed GAIN, a lightweight model that implements these principles by predicting the pseudobulk response to a combination as the sum of its two single-perturbation responses and a learned interaction term. Across three single-cell datasets of genetic and drug combinations, GAIN corrected deviations from additivity and out-performed the additive model and cell-level methods across generalization settings and metrics, while training 3-to 64-fold faster. Applied to a combinatorial CRISPR screen of leukemia differentiation regulators, GAIN revealed synergy between key regulators of the differentiation block and genes that restrain differentiation but have little effect alone. It also revealed dampening by the histone chaperone FACT, which supports the induction of differentiation genes. These results establish population-level modeling of the deviation from additivity as a simple, accurate and scalable approach to predicting combination responses.

## 1 Introduction

Combinatorial perturbations are widely used in both basic and translational biology. Genetic interactions reveal how genes cooperate within pathways, and drug combinations are widely used to increase efficacy and overcome resistance [1, 2]. Advances in single-cell perturbation screening now allow transcriptional responses to combinations to be measured at scale, for both genetic and chemical perturbations [3–7]. However, the number of possible combinations grows quadratically with the number of perturbations, and only a small fraction can be tested experimentally in any given cellular context. Computational methods that predict responses to unmeasured combinations from measured ones are therefore needed.

A variety of deep-learning models have been proposed for this task. Most of these models represent each perturbation as a feature vector and train a neural network to map control cells to perturbed cells. The perturbation features are commonly derived from prior knowledge, such as gene–gene graphs built from functional annotations, or from molecular structure in the case of drugs [8, 9]. Some models further learn the full distribution of perturbed cells within generative frameworks, such as latent-variable models or flow matching [6, 10, 11]. Another class of methods leverages foundation models pretrained on large collections of single-cell transcriptomes or biological text. These methods either use the learned gene embeddings as perturbation features or simulate perturbations in silico [12–15]. Despite this growing model complexity, there is ongoing debate about whether these models outperform simple linear baselines, such as the additive model, which predicts the response to a combination as the sum of its single-perturbation responses. The answer depends strongly on the evaluation metric, and it also varies across datasets and generalization settings [16–19].

We note two aspects of the current approaches that may limit their advantage over additive baselines. First, most of these models predict expression at the single-cell level, whereas their predictions are evaluated at the population level. Because single-cell measurements are sparse and noisy, and the same cell cannot be profiled simultaneously before and after perturbation, perturbation responses can only be quantified by comparing cell populations, for example as changes in mean expression or in expression distributions between conditions. Downstream analyses, such as the identification of responsive genes, are also typically performed at the population level (Fig. 1a). Predicted cells are therefore aggregated before evaluation, and modeling cell-to-cell variation may add complexity without improving the evaluated quantity. Modeling perturbation effects directly at the population level has recently been shown to improve generalization for single perturbations [20]. Second, existing models typically learn the response to a combination directly from control expression data, without using the measured responses to its constituent perturbations. However, at the population level, these single-perturbation responses already explain much of the combination response. In the three single-cell datasets analyzed in this study, the additive model explained a median of 80%, 73% and 47% of the response per combination in the Norman, ComboSciPlex and D-SPIN datasets, respectively (Fig. 1b), consistent with the strong performance of additive baselines reported previously [16]. Measured single-perturbation responses therefore provide an informative prior, leaving the model to predict only the part of the combination response that their sum does not explain.

**Fig 1.**
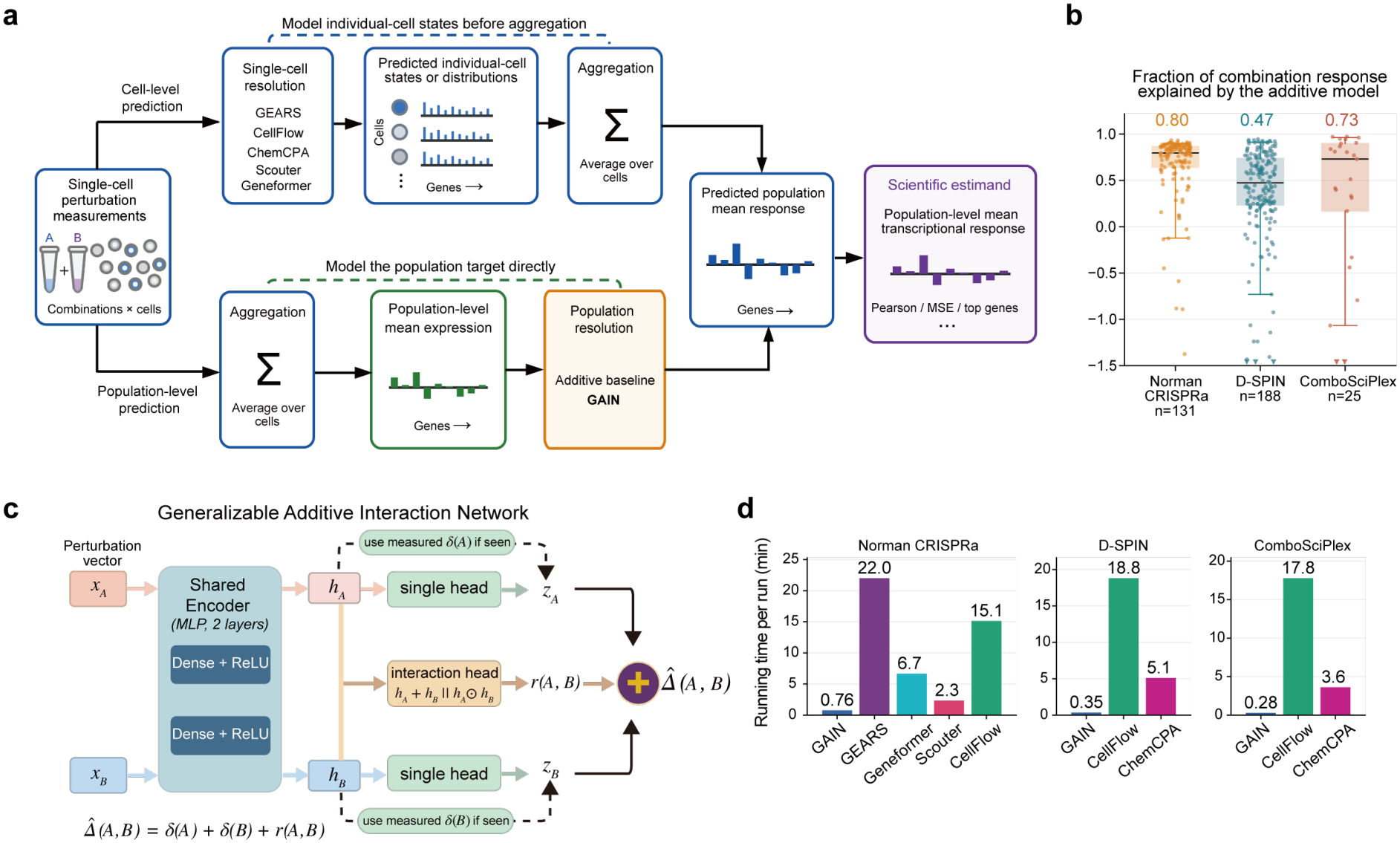
Population-level prediction of combination responses with GAIN. **a**, The schematic shows two routes from single-cell perturbation measurements to the population-level mean response. Cell-level models, namely GEARS, CellFlow, ChemCPA, Scouter and Geneformer, predict individual-cell states or distributions, which are then averaged over cells. The additive model and GAIN predict the population-level mean response directly from pseudobulk profiles. Both routes are evaluated against the same measured population-level response. **b**, Box plots show the fraction of each combination response explained by the additive model for the seen2 test combinations, in which both single-perturbation responses are measured. This fraction is defined as 1 *− ∥ Δ*(*A, B*) *− δ*(*A*) *− δ*(*B*) *∥*^2^*/ ∥Δ*(*A, B*) *∥*^2^, where *Δ*(*A, B*) is the measured response to the combination of perturbations *A* and *B, δ*(*A*) and *δ*(*B*) are the measured responses to *A* and *B* alone, and *∥·∥* is the Euclidean norm over genes. Each response is the change in mean expression relative to control cells. Each dot is one combination, and for D-SPIN each cell type is counted separately. Boxes show the quartiles and whiskers the 5th and 95th percentiles. Numbers above the boxes are medians. **c**, The schematic shows the architecture of GAIN. A shared encoder maps the features *x*_*A*_ and *x*_*B*_ of the two perturbations to embeddings *h*_*A*_ and *h*_*B*_ . A single-perturbation head predicts the response to each perturbation, and the measured response is used instead when it is available. An interaction head predicts the interaction term *r*(*A, B*) from the sum *h*_*A*_ + *h*_*B*_ and the elementwise product *h*_*A*_ *⊙ h*_*B*_ of the two embeddings. The predicted combination response is the sum of the two single-perturbation responses and the interaction term. **d**, The bar plot shows the mean training time of each model per run, where a run is one fold of one setting.

Here, we developed GAIN (Generalizable Additive Interaction Network; Fig. 1c), a lightweight model for predicting combinatorial perturbation responses, to test whether combining population-level prediction with this additive prior is advantageous. GAIN operates on pseudobulk profiles, obtained by averaging expression over cells that received the same perturbation. GAIN predicts the response to the combination of perturbations A and B as the sum of three terms. The first two are the responses to A and to B alone. When a single perturbation has been measured, its observed response is used directly; otherwise, the response is predicted from perturbation features, such as gene features for genetic perturbations or molecular fingerprints for drugs. The third term is an interaction term that captures the deviation of the combination response from the sum of the single responses. It is predicted by a small network from the encoded features of A and B and is symmetric in the two perturbations. Because GAIN does not model individual cells, it trains 3-to 64-fold faster than the cell-level models evaluated here (Fig. 1d).

Across three single-cell datasets of genetic and chemical combinations, GAIN achieved the best or second-best performance among all models, including the additive baseline, deep-learning models and foundation-model-based methods, in 60 of 63 comparisons across datasets, generalization settings and evaluation metrics, and the best in 48 of them. We found that replacing the perturbation features in GAIN with more sophisticated embeddings (e.g., knowledge-informed gene embeddings) or adding a graph encoder did not further improve performance. We then applied GAIN to a combinatorial CRISPR screen of regulators of myeloid differentiation in leukemia cells, in which fewer than half of the possible gene pairs had been measured. GAIN predicted the direction of interaction on the differentiation marker CD11b for 77% of the held-out pairs, compared with 53% for the additive model. Its predictions for the 167 untested pairs completed the map of pairwise interactions. Among them, the strongest predicted synergies combined a key regulator of the differentiation block with a gene that has little effect alone but restrains myeloid differentiation, and the strongest predicted dampening involved the histone chaperone FACT, which supports the induction of differentiation genes. These results suggest that modeling the deviation from additivity at the population level is a simple and effective alternative to modeling individual cells for predicting combination responses. Because this lightweight model is also several-fold faster to train than cell-level models, it makes accurate prediction of combination responses practical for large combinatorial screens.

## 2 Methods

GAIN predicts the response to a combination as the sum of the single-perturbation responses and an interaction term. Each dataset contains control cells, cells treated with a single perturbation and cells treated with a combination of two perturbations. For a single perturbation *A* and the combination of *A* and *B*, we define the single-perturbation response *δ*(*A*) *∈* ℝ ^*G*^ and the combination response *Δ*(*A, B*) *∈* ℝ^*G*^ as

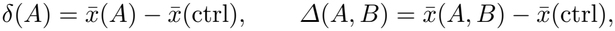

where 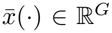 is the log-normalized pseudobulk profile of a condition, obtained by aggregating its cells, and *G* is the number of genes.

GAIN predicts responses in a *d*-dimensional subspace that captures most of the variation among the training responses, with *d ≪ G*. That is, with *V ∈* ℝ^*d×G*^ an orthonormal basis of this subspace computed from the training set only, the measured responses are projected as *z*_*A*_ = *δ*(*A*)*V* ^*⊤*^ and *z*_*AB*_ = *Δ*(*A, B*)*V* ^*⊤*^. GAIN predicts the projected combination response as

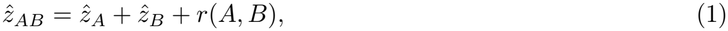

where 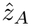 and 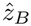 are single-perturbation terms and *r*(*A, B*) is an interaction term. A single-perturbation term equals the measured projected response when that response is in the training set and is otherwise predicted from the features of the perturbation,

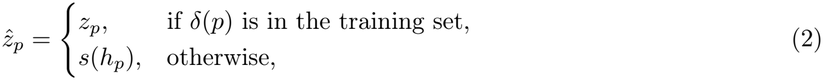

and the interaction term is predicted from both perturbations,

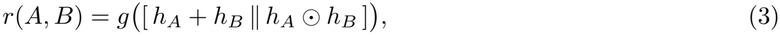

where *h*_*p*_ = *f* (*x*_*p*_) is the latent representation of perturbation *p*, computed by a shared encoder *f* from the perturbation vector *x*_*p*_ of *p*; *s* and *g* are neural networks; *∥* denotes concatenation; and *⊙* denotes elementwise multiplication. For a genetic perturbation, *x*_*p*_ is the PCA embedding of the perturbed gene computed from control cells, and for a drug, it is the Morgan fingerprint of the molecule [21]. The last layer of *g* is initialized to zero, so training starts from the additive prediction 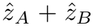. The prediction is mapped back to gene space as 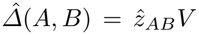. The architecture and training procedure are described in the Supplementary Information.

We evaluated GAIN on three single-cell datasets, Norman CRISPRa with genetic combinations [4], and D-SPIN [7] and ComboSciPlex [6] with drug combinations. In each dataset, the combinations were split into a training set and a test set. Test combinations were split into seen2, seen1 and seen0 by whether two, one or none of their single-perturbation responses were in the training set. We compared GAIN with the additive model, which predicts *δ*(*A*) + *δ*(*B*) and sets the response of an unmeasured perturbation to zero, and with models that predict expression in individual cells, including GEARS [8], Scouter [15], Geneformer [12] and CellFlow [11] for genetic perturbations, and CellFlow and ChemCPA [9] for drug perturbations (Supplementary Fig. 1). Prediction quality was measured for each test combination with metrics including the Pearson correlation, the overlap of the top 20 response genes (top-response-gene overlap) and the calibrated metrics WMSE, weighted 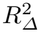 and NIR [18, 19]. We also applied GAIN to a combinatorial CRISPR screen with transcriptome and surface protein measurements [5].

## 3 Results

### 3.1 GAIN outperforms the additive model where cell-level models do not

We benchmarked GAIN against the additive model and the cell-level models across the three datasets, three settings and seven metrics. GAIN ranked first in 48 and second in 12 of 63 comparisons (Fig. 2a; Supplementary Fig. 2). The metrics capture two aspects of performance. The Pearson correlation, top-response-gene overlap and direction accuracy evaluate the pattern of the response, whereas the mean squared error (MSE), weighted mean squared error (WMSE), weighted coefficient of determination (weighted 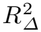) and normalized inverse rank (NIR) also account for its magnitude, and the last three are calibrated metrics that focus on the response specific to each perturbation [18, 19] (Supplementary Information). In seen2, the additive model was a strong baseline, with a Pearson correlation of 0.91 on Norman. The cell-level models fell below it on most metrics. GAIN ranked first in 16 of 21 comparisons and second in 4, with the largest gains in response magnitude. On D-SPIN, for example, it reduced the WMSE of the additive model from 0.051 to 0.017. In seen1 and seen0, at least one single-perturbation response is not measured and must be predicted from perturbation features. GAIN remained the leading model in these settings, ranking first in 32 and second in 8 of 42 comparisons. In seen1, it ranked first in 19 of 21 comparisons and second in 2. Its advantage over the additive model was larger than in seen2, because the additive model sets unmeasured responses to zero (Pearson correlation 0.81 versus 0.61 on D-SPIN). In seen0, GAIN ranked first on every metric on D-SPIN and on all metrics but direction accuracy on ComboSciPlex. Its Pearson correlations of 0.69 and 0.67 on these two datasets compared with at most 0.50 and 0.51 for the cell-level models. On Norman, GAIN and CellFlow performed similarly (0.54 and 0.55), and GAIN ranked second on five of the seven metrics, behind CellFlow or GEARS. On Norman in seen2, the NIR of the leading models was close to its maximum of 1 and did not distinguish models. In seen0, all models had low NIR and weighted 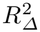, and their differences were smaller than the variation across folds. Moreover, D-SPIN profiles the same drug combinations in four immune cell types, which respond differently to the same drugs [7]. GAIN had the highest Pearson correlation, top-response-gene overlap and weighted 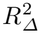 and the lowest MSE and WMSE in every cell type and setting (Supplementary Fig. 3).

**Fig 2.**
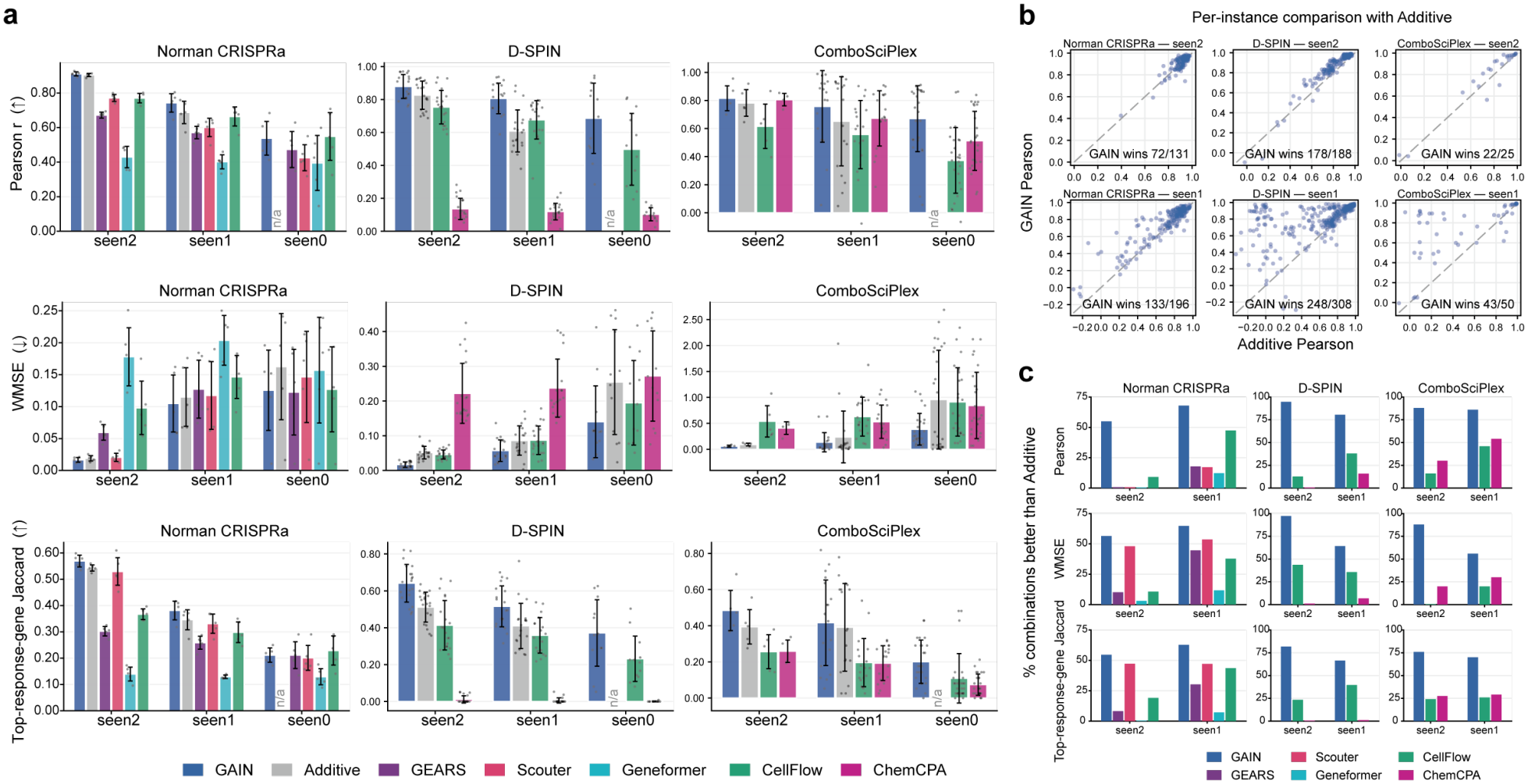
Benchmark of GAIN against the additive model and cell-level models. **a**, Bar plots show the Pearson correlation, WMSE and top-response-gene overlap of each model on Norman, D-SPIN and ComboSciPlex in seen2, seen1 and seen0, in which both, one or neither of the two single-perturbation responses are in the training set. WMSE is the mean squared error with genes weighted by how specifically they respond to the combination. The top-response-gene overlap is the Jaccard index between the predicted and measured sets of the 20 genes with the largest absolute change. Arrows indicate favorable directions. Bars show the mean and error bars the standard deviation across folds, and dots show individual folds. For D-SPIN, each cell type and fold is counted separately. In seen0, the additive prediction is zero, so its Pearson correlation and top-response-gene overlap are undefined and marked n/a. **b**, Scatter plots show the Pearson correlation of GAIN against that of the additive model for each test combination in seen2 and seen1, with each D-SPIN cell type counted separately. Numbers give the combinations on which GAIN had the higher Pearson correlation. **c**, Bar plots show the percentage of test combinations on which each model outperformed the additive model, for the Pearson correlation, WMSE and top-response-gene overlap. Ties count as half.

The advantage of GAIN was not driven by a few combinations. In seen2, GAIN had a higher Pearson correlation than the additive model for 55% of combinations on Norman, 88% on ComboSciPlex and 95% on D-SPIN (Fig. 2b). It outperformed the additive model on more than half of the combinations for every dataset and metric in seen2 and seen1 (Fig. 2c), whereas the cell-level models rarely did. In seen2, the cell-level models outperformed the additive model on fewer than 25% of combinations in 19 of 24 model– dataset–metric comparisons. We next compared the distributions of per-combination scores and the predicted responses themselves. The per-combination scores of GAIN were shifted as a whole relative to those of the other models (Supplementary Fig. 4). In the space of the two leading principal components of the measured and predicted responses, GAIN predictions lay closest to the measured response of each combination (Supplementary Fig. 5). ChemCPA predicted nearly the same response for every combination on D-SPIN, CellFlow predictions were shrunk toward the center, and the additive predictions overshot the measured responses. On ComboSciPlex, ChemCPA had a Pearson correlation close to that of GAIN (0.81 versus 0.82), but its WMSE was nearly seven times that of GAIN. The fraction of combinations improved by GAIN followed the additivity of each dataset (Fig. 1b). It was lowest on Norman, where the additive model explained a median of 80% of the response, and highest on D-SPIN, where it explained 47%. Learning the interaction term therefore helped most where the single-perturbation responses explained least of the combination response.

### 3.2 GAIN captures sub-additive interactions that the additive model misses

We grouped the Norman combinations by the five interaction classes defined by Norman et al.: synergy, redundancy, neomorphism, epistasis and suppression [4] (Fig. 3a,b; Supplementary Fig. 6a). Across classes and metrics, GAIN ranked first in 22 and second in 12 of 35 comparisons in seen2, and first in 25 and second in 6 in seen1. GAIN improved on the additive model mainly in the classes where the additive model was less accurate, namely synergy, neomorphism and suppression. In particular, for suppression, it ranked first on all seven metrics in both settings. For redundancy and epistasis, the two models predicted the pattern of the response similarly, because the combination response resembles one or both single-perturbation responses in these classes [4] and the additive model was already accurate (Pearson correlation 0.95 and 0.93). For redundancy, however, GAIN corrected the magnitude of the response. It reduced the WMSE from 0.043 to 0.029 and increased the weighted 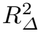 from 0.76 to 0.85 in seen2. In seen1, GAIN improved on the additive model in every class except redundancy, most strongly for epistasis, with a Pearson correlation of 0.63 compared with 0.47. In contrast, the cell-level models rarely outperformed the additive model in seen2. The best of them, Scouter, did so in 8 of 35 comparisons. These cell-level models predict the expression after the combined perturbation de novo from control cells, without making full use of the measured single-perturbation responses.

**Fig 3.**
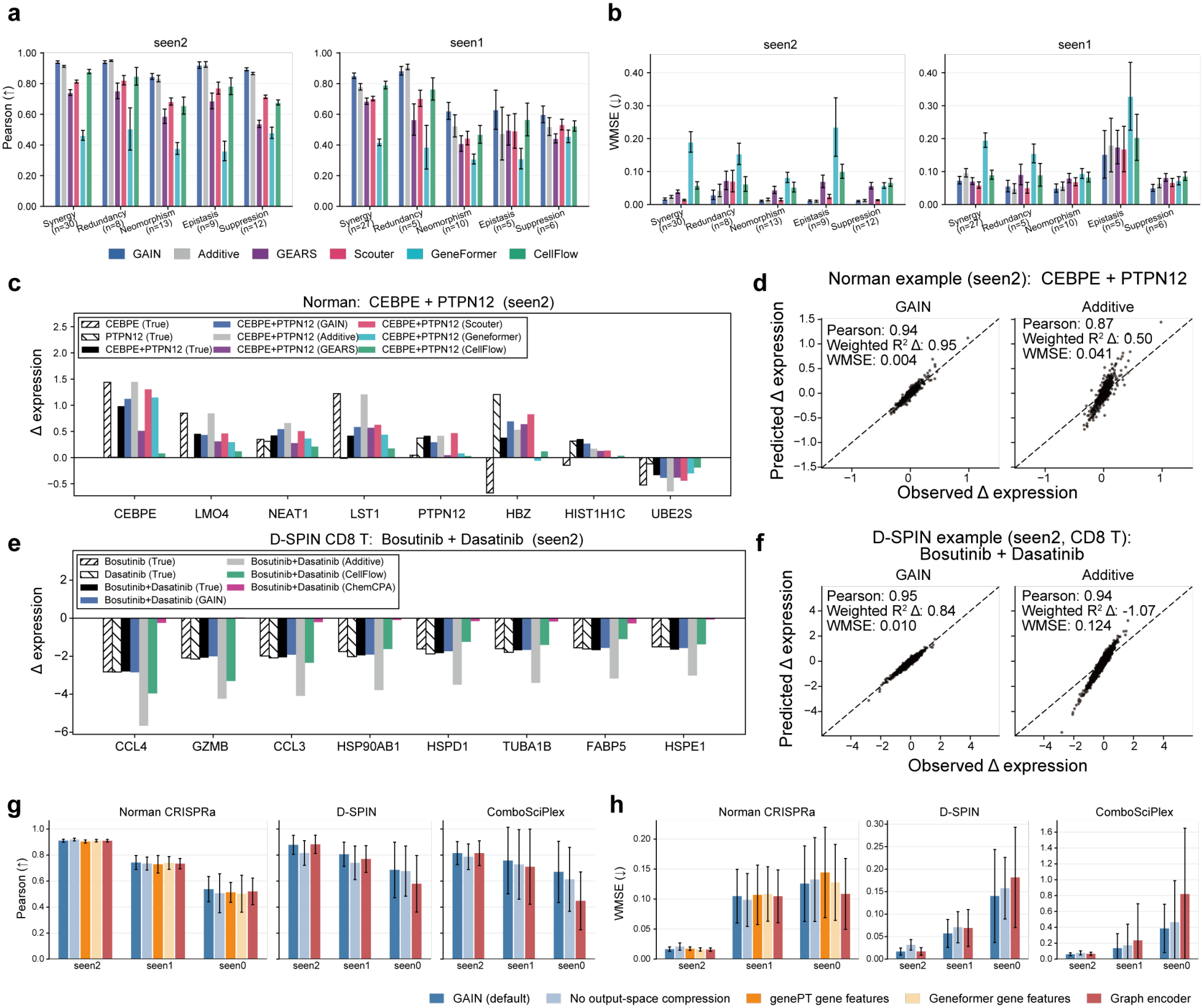
GAIN captures sub-additive interactions that the additive model misses. **a**,**b**, Bar plots show the Pearson correlation (**a**) and WMSE (**b**) of each model on Norman combinations grouped by the five interaction classes defined by Norman et al. [4], in seen2 and seen1. Arrows indicate favorable directions. Bars show the mean and error bars the standard error of the mean across combinations, and *n* is the number of combinations in each class. **c**, Bar plots show the measured and predicted responses to CEBPE + PTPN12 in Norman (seen2) for the eight genes with the largest measured change in the combination. Hatched bars show the measured single-perturbation responses. **d**, Scatter plots show the predicted against the observed responses of all genes to CEBPE + PTPN12 for GAIN and the additive model. Each dot is one gene, and values give the Pearson correlation, weighted 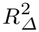 and WMSE of each prediction. **e**, Bar plots show the measured and predicted responses to Bosutinib + Dasatinib in D-SPIN CD8 T cells (seen2) for the eight genes with the largest measured change in the combination. Hatched bars show the measured single-drug responses. **f**, Scatter plots show the predicted against the observed responses of all genes to Bosutinib + Dasatinib for GAIN and the additive model. Each dot is one gene, and values give the Pearson correlation, weighted 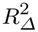 and WMSE of each prediction. **g**,**h**, Bar plots show the Pearson correlation (**g**) and WMSE (**h**) of GAIN and its variants on Norman, D-SPIN and ComboSciPlex in the three settings. The GenePT and Geneformer variants replace the control-expression gene features with pretrained gene embeddings and apply only to Norman. The graph-encoder variant adds a graph of similar perturbations to the perturbation encoder. The variant without output-space compression predicts responses in the full gene space instead of the subspace of training responses. Arrows indicate favorable directions. Bars show the mean and error bars the standard deviation across folds. For D-SPIN, each cell type and fold is counted separately.

We next examined the predictions for one combination from each of the Norman and D-SPIN datasets whose response was much smaller than the sum of the single-perturbation responses, that is, a sub-additive combination. Based only on the measured responses, we selected the combination with the highest suppression ratio, which is defined as the magnitude of the summed single-perturbation responses divided by that of the combination response (Supplementary Information). In Norman, CEBPE + PTPN12 was selected with the highest suppression ratio among suppression combinations, 2.04 across all genes. GAIN predicted this response more consistently with the measured values than the other models (Fig. 3c). On the eight genes with the largest measured change, the mean absolute error was 0.13 for GAIN, 0.18 for Scouter, 0.20 for Geneformer, 0.23 for GEARS, 0.32 for the additive model and 0.35 for CellFlow. Specifically, the additive model had its largest errors in the response to CEBPE. Activation of CEBPE alone induced a strong response, but this response was weaker when PTPN12 was also activated. The additive model retained only the full effect of CEBPE, whereas GAIN predicted the reduced response through the interaction term. Across all genes, GAIN achieved the highest Pearson correlation and weighted 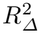 and the lowest WMSE of all models (Fig. 3d; Supplementary Fig. 6b).

We performed the same analysis with a redundant combination on D-SPIN. In a redundant drug pair, the two single-drug responses are similar and the combination response is close to either of them. Bosutinib + Dasatinib in CD8 T cells was selected with similar single-drug responses and the highest suppression ratio, 1.89 (Fig. 3e). Both drugs are known to inhibit SRC and ABL kinases [22, 23]. Because the combination response was close to either single-drug response, the additive model predicted about twice the measured change. On the eight genes with the largest measured change, the mean absolute error was 0.08 for GAIN, 0.59 for CellFlow, 1.82 for ChemCPA, which predicted almost no change, and 1.89 for the additive model. Across all genes, GAIN and the additive model had similar Pearson correlations (0.95 and 0.94), but the weighted 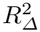 was 0.84 for GAIN and *−*1.07 for the additive model (Fig. 3f; Supplementary Fig. 6c). This indicates that GAIN corrected the magnitude of the response, which the Pearson correlation does not capture. In both examples, the additive model overestimated the magnitude of the sub-additive response, and GAIN corrected this error.

Finally, we asked whether richer perturbation embeddings improve GAIN (Fig. 3g,h). Foundation models and deep-learning models represent perturbations with pretrained embeddings or with graphs that connect related perturbations [8, 12]. We replaced the control-expression gene features of GAIN with GenePT or Geneformer gene embeddings [12, 14], or added a graph encoder that aggregates the features of similar perturbations. None of these variants improved on GAIN. On Norman, all variants performed similarly. In seen1, for example, the Pearson correlation was 0.74 for GAIN and between 0.73 and 0.74 for the variants. On the drug datasets, the graph encoder lowered performance in seen1 and seen0, for example from 0.69 to 0.58 on D-SPIN in seen0. These results suggest that complex perturbation embeddings are not needed for a lightweight model such as GAIN.

### 3.3 GAIN completes the unmeasured gene pairs of a leukemia differentiation screen

We next applied GAIN to the combinatorial CRISPR screen of Wessels et al. [5]. In the acute myeloid leukemia cell line THP-1, this study used Cas13 to knock down one or two of 26 regulators of myeloid differentiation, and measured the transcriptome and 22 surface proteins in the same cells. The original study used the surface protein CD11b as the readout of myeloid maturation; in the transcriptome, this marker corresponds to ITGAM, the gene encoding CD11b. Of the 325 possible pairs of the 26 genes, only 158 were measured. We trained GAIN separately on the transcriptome and on the surface proteins of all measured perturbations and predicted the responses of the remaining 167 pairs, completing the map of all pairwise combinations (Fig. 4a, shown for CD11b protein and ITGAM; dotted squares are GAIN predictions). We first assessed the accuracy of GAIN on the measured pairs and then analyzed the completed pairs.

**Fig 4.**
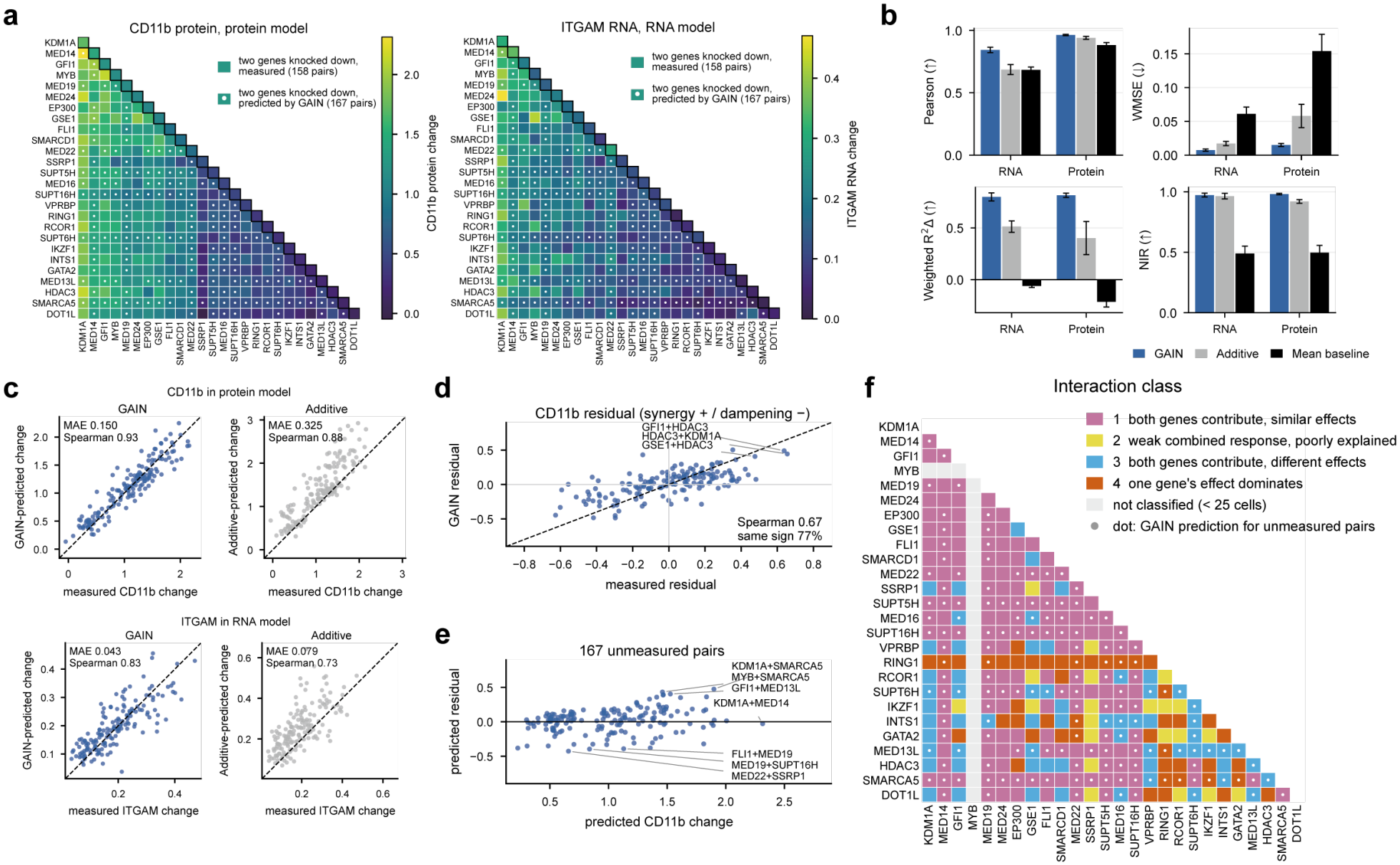
GAIN completes the unmeasured gene pairs of a leukemia differentiation screen. **a**, Heatmaps show the change in CD11b protein and in ITGAM RNA relative to non-targeting control cells for all pairs of the 26 regulators. Unmeasured pairs were predicted by GAIN trained on the surface proteins for CD11b and on the transcriptome for ITGAM separately. The diagonal shows single-gene knockdowns and the lower triangle shows pairs. Squares with a white dot are the 167 unmeasured pairs predicted by GAIN, and the other squares are the measured responses of the 158 measured pairs. **b**, Bar plots show the Pearson correlation, WMSE, weighted 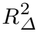 and NIR of GAIN, the additive model and a mean baseline on held-out measured pairs (seen2) for the transcriptome (RNA) and the 22 surface proteins (Protein). The mean baseline predicts the mean response of the training pairs. Arrows indicate favorable directions. Bars show the mean and error bars the standard deviation across five folds. **c**, Scatter plots show the predicted against the measured change of CD11b in the protein model and of ITGAM in the RNA model for held-out pairs, for GAIN and the additive model. Each dot is one held-out pair, and values give the mean absolute error (MAE) and the Spearman correlation. **d**, The scatter plot shows the CD11b residual predicted by GAIN against the measured residual for each held-out pair. The residual of a pair is its deviation from the average linear relationship between the CD11b change of a pair and the mean CD11b change of its two single-gene knockdowns [5]. Positive values indicate synergy and negative values dampening. Values give the Spearman correlation and the percentage of pairs whose predicted and measured residuals have the same sign. The three synergistic pairs reported in the original study are labeled. **e**, The scatter plot shows the predicted residual against the predicted CD11b change for the 167 unmeasured pairs. Pairs with the largest positive and negative residuals are labeled. **f**, The heatmap shows the interaction class of each pair. Classes were defined by clustering the regression features of the measured pairs. Unmeasured pairs, marked with a dot, were assigned to the nearest class from the GAIN predictions. Pairs with fewer than 25 cells for either knockdown were not classified.

GAIN accurately predicted the responses of held-out measured pairs in the seen2 setting, on both the transcriptome (9,391 genes) and the 22 surface proteins. It outperformed the additive model and the mean baseline on all four metrics for both readouts (Fig. 4b). On the transcriptome, GAIN had a Pearson correlation of 0.84 and a weighted 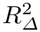 of 0.80, compared with 0.69 and 0.51 for the additive model. On the proteins, these values were 0.96 and 0.82 for GAIN, compared with 0.94 and 0.40 for the additive model. GAIN also accurately predicted the change in CD11b. The predicted changes of the held-out pairs relative to non-targeting control cells closely followed the measured changes (Fig. 4c), with a mean absolute error of 0.15, half that of the additive model (0.33). With transcriptomic data, GAIN predictions of ITGAM were similarly accurate (mean absolute error 0.043 versus 0.079; Fig. 4c).

Beyond the magnitude of the responses, GAIN also captured which pairs acted synergistically (Fig. 4d). We measured the interaction of each pair by its deviation from the average linear relationship between the CD11b change of a pair and the mean CD11b change of its two genes knocked down alone, as originally defined by Wessels et al. [5, Fig. 4c]. A positive deviation indicates synergy and a negative deviation indicates dampening. GAIN predicted the correct direction for 121 of the 158 held-out pairs (77%), whereas the additive model was close to chance (53%), and the predicted deviations correlated with the measured ones (Spearman correlation 0.67 versus 0.06 for the additive model). All three synergistic pairs reported in the original study, HDAC3+KDM1A, GFI1+HDAC3 and GSE1+HDAC3, were predicted as synergistic. GAIN trained on the transcriptome similarly predicted the direction of interaction on ITGAM for 74% of the held-out pairs, compared with 46% for the additive model (Supplementary Fig. 7a).

We then extended this analysis to the 167 unmeasured pairs predicted by GAIN, using the relationship fitted on all measured pairs (Fig. 4e; Supplementary Fig. 7b for ITGAM). The unmeasured pair with the largest predicted increase in CD11b was KDM1A+MED14 (2.31), comparable to the largest measured increase, KDM1A+MED24 (2.16). However, its deviation was close to zero, because KDM1A and MED14 had the two strongest single-gene effects on CD11b among the 26 genes, for which the fitted relationship already predicts an increase of 2.34. The large effect of KDM1A+MED14 therefore arose from two strong single-gene effects rather than from synergy. The pairs with the strongest predicted synergy were KDM1A+SMARCA5, MYB+SMARCA5 and GFI1+MED13L. In each of these pairs, one gene strongly increased CD11b when knocked down alone, whereas the other had almost no effect (0.15 for SMARCA5 and 0.22 for MED13L). This pattern matches the synergistic pairs reported in the original study: knockdown of HDAC3 alone increased CD11b by only 0.21, whereas its combinations with KDM1A, GFI1 or GSE1 exceeded the sum of the two single-gene effects. The weak partners are biologically plausible amplifiers of differentiation. SMARCA5 cooperates with CTCF to repress SPI1 [24], which encodes PU.1, the myeloid transcription factor that activates the CD11b promoter [25] and is required for the differentiation induced by LSD1 inhibition [26]. MED13L belongs to the kinase module of Mediator, whose inhibition further activates myeloid lineage genes in acute myeloid leukemia [27]. Knocking down either gene may thus have little effect while the differentiation block is intact, but amplify differentiation once it is released by knocking down KDM1A, GFI1 or MYB. Conversely, 8 of the 10 pairs with the strongest predicted dampening, including the top two, contained SSRP1 or SUPT16H, which together form the histone chaperone FACT, although such pairs made up only 17% of the unmeasured pairs. FACT allows RNA polymerase II to transcribe through nucleosomes [28] and is recruited to lineage-specific genes as differentiation begins, as shown in muscle cells [29], so its knockdown may blunt the differentiation triggered by the other gene. GAIN also predicted strong dampening for MED19 combined with the transcription factors FLI1 or MYB, consistent with the dependence of the MYB-driven program in acute myeloid leukemia on Mediator [30]. The completed map thus nominates synergistic and dampening pairs that were not tested in the original screen as candidates for experimental validation.

Beyond CD11b, we further characterized the interactions of the unmeasured pairs across the whole transcriptome. Following the original study [4, 5], we fitted the expression change of each pair as a weighted sum of the changes of its two genes knocked down alone, and used six features computed from the fitted coefficients and the goodness of fit to group the measured pairs into four classes (Supplementary Fig. 7c,d). In class 1, both genes contribute equally and have similar effects. In class 2, the combined response is weak and is not explained by the two single-gene responses. In class 3, both genes contribute equally but have different effects. In class 4, the effect of one gene dominates. The three synergistic pairs reported in the original study belonged to class 3. For the unmeasured pairs, we computed the same features from the GAIN predictions and assigned each pair to the nearest class (Fig. 4f). Of the 158 unmeasured pairs that could be classified, 111 belonged to class 1 (Supplementary Fig. 7e), as did all 13 classifiable measured pairs within the same complex. All 9 unmeasured pairs containing RING1 belonged to class 4, as did 12 of the 15 measured pairs containing RING1, consistent with the original study, in which the response to KDM1A knockdown dominated when it was combined with RING1 knockdown [5]. Together, these results show that GAIN can complete a partially measured combinatorial screen: trained on fewer than half of the possible pairs, it recovered the interactions reported in the original study and provided predictions for every untested pair at the levels of the differentiation marker and the whole transcriptome, which can guide the choice of combinations to measure next.

## 4 Discussion

In this study, we proposed that predicting combination responses at the population level, with the measured single-perturbation responses as a prior, is an effective strategy. We implemented this principle with a lightweight model, GAIN, which predicts only how the response to a combination deviates from the sum of its single-perturbation responses. It outperformed the additive model and cell-level deep-learning and foundation-model-based methods across three single-cell datasets of genetic and drug combinations, while training 3-to 64-fold faster. GAIN applies to both transcriptome and surface protein measurements, as shown in the leukemia screen, and could be applied to other readouts. The speed and accuracy of GAIN make it practical for partially measured combinatorial screens. In the leukemia screen, it completed the map of pairwise interactions from fewer than half of the possible pairs, and its predictions can guide the choice of combinations to measure next.

Modeling perturbation responses at the population level or at the level of individual cells involves a trade-off between efficiency and resolution. In our benchmark, the population-level model was more accurate in predicting the mean response and several-fold faster to train, and it directly yields the quantities used in downstream analyses, such as responsive genes and changes in marker expression. However, a mean response cannot distinguish a uniform shift of all cells from a strong response of a subpopulation. Population-level modeling is therefore well suited when the question concerns the average response, such as screening combinations for their effect on a marker or a gene program, and when cells respond relatively uniformly. Cell-level modeling is needed when the distribution of responses itself matters, for example when a combination drives cells into distinct states or eliminates a resistant subpopulation. Intermediate strategies are also possible. Modeling the pseudobulk profiles of cell subsets separately, as we did for the cell types in the D-SPIN dataset, retains part of this heterogeneity, although heterogeneity of cells remains within each subset. In addition, population-level predictions could serve as a prior for cell-level models. A separate limitation of GAIN is that the response to a single perturbation that has not been measured is predicted from perturbation features, and such responses remained difficult to predict when neither perturbation was measured. A dedicated model for single perturbations could replace this component of GAIN. The additive formulation of GAIN extends naturally beyond pairs. The response to a combination of three or more perturbations could be modeled as the sum of the single-perturbation responses and interaction terms of increasing order, and higher-order combinations are even less covered by experiments than pairs. GAIN could also incorporate covariates of the experimental condition, such as the dose and treatment duration of a drug, to predict responses across conditions.

## Code availability

The source code of GAIN is available at https://github.com/caokai1073/GAIN. GAIN can also be used through a web server at https://www.combo-gain.com, where users can upload their own perturbation data and obtain predicted responses to combinations.

## Data availability

Gene-perturbation data were obtained from Norman et al. (2019; GEO: GSE133344) [4] using the harmonized scPerturb AnnData (.h5ad) release (Zenodo: 13350497) [31]. D-SPIN MULT-18 single-cell counts were obtained from Jiang et al. (2026) [7] through CaltechDATA (https://data.caltech.edu/records/2cjss-wgh69). ComboSciPlex data were obtained from Lotfollahi et al. (2023; GEO: GSE206741) [6], with missing single-drug profiles supplemented from the sci-Plex3 dataset (Srivatsan et al., 2020; GEO: GSE139944) [32] through scPerturb. Single-cell CRISPR perturbation data with surface protein measurements were obtained from Wessels et al. (2023; GEO: GSE213957) [5]. All source datasets are publicly available, and the processed response matrices used for model training will be provided with the code release.

