## supplementary method and figures for "Population-level learning of non-additive effects enables accurate and efficient prediction of combinatorial perturbation responses"

### Supplementary Information

#### 1 Supplementary Methods

##### 1.1 Datasets

We used four datasets of combinatorial perturbations, all profiled by single-cell RNA sequencing (Supplementary Fig. 1).

**Norman CRISPRa.** This dataset contains K562 cells in which one or two genes were activated by CRISPR activation and profiled by single-cell RNA sequencing [1]. We used the version processed by scPerturb [2], which contains 111,445 cells, including 11,855 control cells, across 105 single-gene and 131 two-gene perturbations. Each perturbation was profiled in at least 20 cells. We restricted the genes to the 5,045-gene panel used by GEARS [3], of which 5,041 were present in the data. The interaction class of each combination was taken from the original study [1].

**D-SPIN.** This dataset contains human peripheral blood mononuclear cells treated with single drugs and pairs of drugs under anti-CD3 stimulation and profiled by single-cell RNA sequencing [4]. We used the batch MULT-18, in which the control, 18 single drugs and 65 pairs of these drugs were profiled in one pooled experiment, so that single-drug and combination conditions share the same batch. Because all drug conditions were stimulated, we used the stimulated control cells as the reference. We analyzed the four coarse cell types with sufficient cells, namely B cells, CD4 T cells, CD8 T cells and myeloid cells, and modeled each cell type separately. In each cell type, we kept the conditions profiled in at least 20 cells and the combinations whose two single-drug conditions were both kept. This yielded 14 to 16 single drugs and 36 to 54 combinations per cell type, with 188 combinations and 49,953 cells in total. In each cell type, we kept the genes with at least 50 counts summed over all conditions, which gave 10,945 to 14,265 genes.

**ComboSciPlex.** This dataset contains A549 cells treated with 25 pairs of drugs, each at 1 $\mu$M, for 24 h and profiled by sci-Plex [5]. It also includes single-drug profiles for 6 of the 17 drugs. For the other 11 drugs, we obtained single-drug profiles from sci-Plex3, which profiled A549 cells treated at 10 $\mu$M for 24 h [6]. We restricted the genes to those measured in both datasets and selected the 2,000 most highly variable genes among them in the ComboSciPlex cells with Scanpy [7], after library-size normalization and log transformation.

**Wessels CRISPR knockdown.** This dataset contains THP-1 acute myeloid leukemia cells in which one or two of 28 genes, namely 26 regulators of myeloid differentiation and two negative control genes, were knocked down by Cas13 and profiled by CaRPool-seq, which measures the transcriptome and surface proteins in the same cells [8]. We used the cells that passed quality control and had a guide assignment in the metadata released with the study, which gave 30,707 cells, including 424 control cells with non-targeting guides, across 28 single-gene and 158 two-gene perturbations. The 158 pairs involve 26 of the 28 genes and cover 158 of the 325 possible pairs of these genes. We kept the 9,391 genes detected in at least 5% of the cells. The released antibody matrices list 58 antibodies, but the THP-1 screen was stained with a panel of 22 antibodies [8, Supplementary Table 6], so we kept these 22 surface proteins.

##### 1.2 Data preprocessing

Each dataset contains control samples, samples treated with a single perturbation and samples treated with a combination of two perturbations. For a single perturbation $A$ and the combination of $A$ and $B$, we defined the single-perturbation response $\delta(A) \in \mathbb{R}^G$ and the combination response $\Delta(A, B) \in \mathbb{R}^G$ as

$$\delta(A) = \bar{x}(A) - \bar{x}(\text{ctrl}), \qquad \Delta(A, B) = \bar{x}(A, B) - \bar{x}(\text{ctrl}), \tag{S1}$$

where $G$ is the number of genes, and $\bar{x}(A)$, $\bar{x}(A, B)$ and $\bar{x}(\text{ctrl})$ are the average log-normalized expression profiles of the samples treated with $A$, the samples treated with $A$ and $B$, and the control samples. Depending on the dataset, these profiles were averaged over cells or computed from pseudobulk profiles, as described below. In all datasets, expression was normalized by library size and transformed by $\log(1 + x)$.

**Norman CRISPRa.** We normalized and log-transformed each cell and averaged the cells of each perturbation and of the control, following the standard preprocessing of this dataset [3]. We then restricted the responses to the GEARS gene panel (Datasets).

**D-SPIN and ComboSciPlex.** For each condition, we summed the raw counts of its cells into a pseudobulk profile and then normalized and log-transformed this profile. In D-SPIN, we did this in each cell type separately, normalized over the kept genes and took the mean of the six stimulated control replicates as the control profile. In ComboSciPlex, the responses to combinations and to the six drugs profiled alone were computed relative to the DMSO control of the same dataset, and the responses to the 11 drugs from sci-Plex3 were computed relative to the vehicle control of sci-Plex3. We matched genes between the two datasets by Ensembl identifier and restricted all responses to the 2,000 highly variable genes (Datasets).

**Wessels CRISPR knockdown.** As for Norman CRISPRa, we normalized and log-transformed each cell and averaged the cells of each perturbation and of the non-targeting control. For the surface proteins, we applied the centered log-ratio transformation to each cell over the 22 antibodies, that is, $\log(1+x)$ minus its mean over the 22 antibodies of the cell, and computed the responses in the same way.

These responses were the training and evaluation targets of GAIN and the additive model. Cell-level models were trained on the log-normalized single-cell profiles, and their predicted cells were aggregated in the same way as the measured responses of each dataset (Baseline methods).

##### 1.3 Evaluation settings

We evaluated each method on combinations held out from training. Following GEARS [3], we divided the test combinations into three settings by how many of their single perturbations were in the training set. In seen2, both single perturbations were in the training set, so this setting tests whether a combination response can be predicted from measured single-perturbation responses. In seen1, one of the two single perturbations was in the training set, and in seen0, neither was. All methods were trained and evaluated on the same folds, and in D-SPIN the folds were constructed separately for each cell type.

**seen2.** We randomly divided the combinations whose two single perturbations were both measured into five folds. In each fold, the training set contained all single perturbations and the combinations of the other four folds.

**seen1 and seen0.** For Norman and D-SPIN, we randomly divided the perturbed genes or drugs into five groups and held out one group in each fold. We removed from the training set every condition that contained a held-out perturbation, including its single-perturbation profile and all combinations with it, so that held-out perturbations were never seen during training. Test combinations with exactly one held-out perturbation formed seen1, and those with two held-out perturbations formed seen0. Both settings used the same held-out groups. In D-SPIN, some folds contained no test combinations, which left three folds for seen0 in each cell type and four folds for seen1 in CD4 T cells.

**ComboSciPlex.** Because this dataset contains only 25 combinations, random groups of drugs would leave many folds without test combinations. We therefore held out one drug at a time for seen1, which gave 18 folds, and the two drugs of one combination at a time for seen0, which gave 25 folds with one test combination each. As in the other datasets, the held-out drugs were removed from the training set together with all combinations that contained them.

##### 1.4 GAIN

GAIN predicts the response to a combination of two perturbations as the sum of the two single-perturbation responses and an interaction term. Its input is a feature vector for each perturbation, which a shared encoder maps to a latent representation. Two networks take these representations as input. A single-perturbation network predicts the response to one perturbation, and an interaction network predicts how the response to a combination deviates from the sum of the two single-perturbation responses. Both networks predict responses in a low-dimensional subspace, and the predictions are then mapped back to gene space.

To define this subspace, we computed in each fold a truncated singular value decomposition of the training responses, which include the single-perturbation responses and the combination responses in the training set. The top $d$ right singular vectors form an orthonormal basis $V \in \mathbb{R}^{d\times G}$, with $d$ ranging from 15 to 50 across datasets. Each measured response was projected onto this subspace as $z_A = \delta(A)V^\top$ and $z_{AB} = \varDelta(A,B)V^\top$.

In this subspace, GAIN predicts the projected combination response as

$$\hat{z}_{AB} = \hat{z}_A + \hat{z}_B + r(A,B), \tag{S2}$$

where $\hat{z}_A$ and $\hat{z}_B$ are single-perturbation terms and $r(A,B)$ is the interaction term. A single-perturbation term equals the measured projected response when that response is in the training set and is otherwise predicted by the single-perturbation network $s$,

$$\hat{z}_p = \begin{cases} z_p, & \text{if } \delta(p) \text{ is in the training set,} \\ s(h_p), & \text{otherwise.} \end{cases} \tag{S3}$$

The interaction term is predicted by the interaction network $g$ from both perturbations,

$$r(A, B) = g\big([\, h_A + h_B \,\|\, h_A \odot h_B \,]\big), \tag{S4}$$

where $h_p = f(x_p)$ is the latent representation of perturbation $p$, computed by the shared encoder $f$ from its feature vector $x_p$, $\|$ denotes concatenation and $\odot$ denotes elementwise multiplication. Because the sum and the elementwise product are symmetric in $A$ and $B$, the prediction is invariant to the order of the two perturbations. The predicted combination response is mapped back to gene space as $\hat{\Delta}(A, B) = \hat{z}_{AB} V$, and the predicted response to an unmeasured single perturbation as $\hat{\delta}(p) = s(h_p)V$.

GAIN is trained by comparing these predictions with the measured responses in the subspace. For a single perturbation $p$ in the training set, the prediction is $s(h_p)$ and the target is $z_p$. For a combination in the training set, the prediction is $\hat{z}_{AB}$ and the target is $z_{AB}$. With $\mathcal{S}$ and $\mathcal{C}$ denoting the single perturbations and the combinations in the training set, the loss is the mean squared error over the $d$ dimensions,

$$\mathcal{L} = \sum_{p \in \mathcal{S}} \frac{1}{d} \left\| s(h_p) - z_p \right\|_2^2 + \sum_{(A,B) \in \mathcal{C}} \frac{1}{d} \left\| \hat{z}_{AB} - z_{AB} \right\|_2^2. \tag{S5}$$

Single-perturbation examples therefore train $f$ and $s$, and combination examples train $f$ and $g$, together with $s$ when one of the two single-perturbation responses is absent from the training set.

Because $s$ and $g$ act on features rather than on measured responses, GAIN can predict combinations that involve unmeasured perturbations. For a genetic perturbation, $x_p$ is the loading vector of the perturbed gene on the top 128 principal components of the log-normalized control cells. For a drug, $x_p$ is the Morgan fingerprint of the molecule with radius 4 and 1,024 bits, computed with RDKit [9,10]. Both features depend only on control cells or chemical structure, so they are available for every perturbation.

The encoder $f$, the single-perturbation network $s$ and the interaction network $g$ are multilayer perceptrons shared across all perturbations. The last layer of $g$ is initialized to zero, so training starts from the additive prediction $\hat{z}_A + \hat{z}_B$ and $g$ learns only the deviation from it. We trained GAIN with the Adam optimizer [11] at a learning rate of $10^{-3}$.

##### 1.5 Ablation and alternative perturbation representations

To test whether richer perturbation representations improve GAIN, we replaced its gene features with pre-trained gene embeddings or added a graph encoder that aggregates the features of related perturbations. We also removed the compression of the output space to assess its contribution. Each variant changes one component of GAIN and keeps the training sets, loss function, hyperparameters and evaluation unchanged.

**No output-space compression.** This variant predicts responses in gene space without projecting them onto the subspace of training responses. The networks $s$ and $g$ output vectors of length $G$, and the loss is computed over all $G$ genes. The prediction formula is unchanged, so measured single-perturbation responses are still used directly. We applied this variant to all datasets to test whether the compression to a low-dimensional subspace improves prediction.

**Graph encoder.** This variant replaces the multilayer perceptron encoder $f$ with a relational graph convolutional network [12], which aggregates the representations of related perturbations. In each layer $l$, the representation of perturbation $p$ is updated as

$$h_p^{(l+1)} = \mathrm{ReLU}\Big(W_0^{(l)} h_p^{(l)} + \sum_r W_r^{(l)} \sum_q A_{pq}^{(r)}\, h_q^{(l)}\Big), \tag{S6}$$

where $A^{(r)}$ is the adjacency matrix of relation $r$ and $W_0^{(l)}$ and $W_r^{(l)}$ are learned weight matrices. For genetic perturbations, we used four relations between perturbed genes: the absolute Pearson correlation of their expression in control cells, the Jaccard overlap of their GO and Reactome annotations [13, 14], the cosine similarity of their GenePT embeddings, keeping the 10 most similar genes for each gene [15], and the regulatory links between transcription factors and their targets in NicheNet [16]. Each adjacency matrix was normalized by its row sums. For drugs, we used a single relation that connects two drugs when the Tanimoto similarity of their Morgan fingerprints exceeds 0.15. All relations are computed from prior knowledge, control cells or chemical structure, so they are available for unmeasured perturbations.

**Pretrained gene embeddings.** These two variants replace the gene features $x_p$ of Norman CRISPRa with pretrained gene embeddings, namely the 1,536-dimensional GenePT embedding [15] or the 768-dimensional input token embedding of Geneformer [17]. The embeddings were normalized to unit length, and they were available for 104 of the 105 perturbed genes. The feature vector of the remaining gene was set to zero. Because comparable pretrained embeddings are not available for the drugs in our datasets, these two variants were applied only to Norman CRISPRa.

##### 1.6 Baseline methods

We compared GAIN with the additive model on all three datasets. On Norman CRISPRa, we compared it with GEARS [3], Scouter [18], Geneformer [17] and CellFlow [19]. On D-SPIN and ComboSciPlex, we compared it with CellFlow and ChemCPA [20]. The cell-level models were trained on single-cell profiles normalized by library size and transformed by $\log(1 + x)$, following the preprocessing of their original implementations. All methods were trained and evaluated on the same folds (Evaluation settings), and every prediction was converted to a response relative to the control before scoring.
**Additive model.** The additive model predicts the response to the combination of $A$ and $B$ as $\delta(A) + \delta(B)$, the sum of the two measured single-perturbation responses in the training set. When a single perturbation was held out, as in seen1 and seen0, its response was set to zero. In seen0 the additive prediction is therefore zero for every gene.
**GEARS.** GEARS represents each perturbed gene by a node in a gene ontology graph and predicts the expression profile of a perturbation with a graph neural network [3]. We used the official implementation with a hidden dimension of 64 and a batch size of 32. The response was the predicted mean expression minus the mean expression of the control cells.
**Scouter.** Scouter predicts the expression profile of a perturbation from language-model embeddings of the perturbed genes [18]. We used the official implementation with its default settings and GenePT gene embeddings [15].
**Geneformer.** Geneformer outputs cell embeddings rather than expression profiles, so we followed Ahlmann-Eltze et al. [21] and decoded its embeddings with a linear model. We fine-tuned Geneformer V2 [17, 22] to classify the perturbation labels of the training cells and fit a ridge regression from the embedding of each training cell to the mean expression profile of its condition. We embedded a single perturbation by in silico overexpression in control cells [17] and the combination of $A$ and $B$ as $e(A) + e(B) - e(\text{ctrl})$, where $e$ denotes the mean embedding. The predicted response was the ridge prediction minus the mean expression of the control cells.
**CellFlow.** CellFlow learns a flow-matching model that transports control cells to perturbed cells conditioned on embeddings of the perturbations [19]. We used its default network with 50,000 training iterations and a batch size of 1,024. Cells were represented by the first 50 principal components fit on the training cells. Perturbed genes were embedded with ESM2 [23], and drugs were embedded with Morgan fingerprints of radius 4 and 1,024 bits [9]. On Norman CRISPRa, we used 500 control cells held out from training as source cells and averaged the log-normalized predicted cells. On D-SPIN and ComboSciPlex, we used the training control cells as source cells and aggregated the predicted cells into a pseudobulk profile by averaging their expression in linear scale and then applying $\log(1+x)$, to match the measured responses (Data preprocessing).
**ChemCPA.** ChemCPA predicts the response of each cell to a drug from a latent representation of the cell and an embedding of the drug structure [20]. We used the official implementation in the cpa-tools package with RDKit drug embeddings and trained it on raw counts with a negative binomial loss. We used a latent dimension of 32, encoders and decoders with two layers of 128 units. The predicted cells of each condition were summed into a pseudobulk profile, which was normalized and log-transformed as the measured profiles (Data preprocessing).

##### 1.7 Evaluation metrics

All metrics compare the predicted response $\hat{\Delta} \in \mathbb{R}^G$ of one test combination with its measured response $\Delta \in \mathbb{R}^G$, where $G$ is the number of genes. We computed each metric for every test combination and averaged it over the test combinations of each setting.
**Unweighted metrics.** The Pearson correlation and the mean squared error, $\text{MSE} = \frac{1}{G}\sum_g(\Delta_g - \hat{\Delta}_g)^2$, were computed over all $G$ genes. To assess the genes with the largest responses, we took the 20 genes with the largest $|\hat{\Delta}_g|$ and the 20 genes with the largest $|\Delta_g|$, and computed the top-20 Jaccard index as the size

of their intersection divided by the size of their union. The direction accuracy is the fraction of the 20 genes with the largest $|\varDelta_g|$ for which $\hat{\varDelta}_g$ and $\varDelta_g$ have the same sign. In seen0 the additive prediction is zero for every gene, so its Pearson correlation and top-20 metrics are not defined.

**Weighted metrics.** We also computed two metrics in which genes are weighted. Most genes respond weakly to a perturbation, and unweighted metrics can be dominated by them. Following Mejia et al. [24] and Miller et al. [25], we therefore weighted genes by how strongly they distinguish a combination from the other perturbations. For each combination and gene $g$, we computed a $t$-score $t_g$ between the cells of this combination and all other perturbed cells of the dataset, excluding control cells, with the `t-test_overestim_var` method of Scanpy [7]. In D-SPIN, the other perturbed cells were taken from the same cell type. We scaled $|t_g|$ to $[0, 1]$ by min-max normalization, squared it and divided by the sum over genes, so that the weights $w_g$ sum to 1.

Given these weights, the weighted mean squared error is WMSE $= \sum_g w_g(\varDelta_g - \hat{\varDelta}_g)^2$. The weighted $R^2$ of the response, $R^2_w(\varDelta)$, measures how much of the response specific to a combination is predicted. In each fold, we took the mean response $r \in \mathbb{R}^G$ of all training perturbations, including single perturbations and combinations, and subtracted it from both the measured and the predicted responses, which gave $\varDelta' = \varDelta - r$ and $\hat{\varDelta}' = \hat{\varDelta} - r$. Then

$$R^2_w(\varDelta) = 1 - \frac{\sum_g w_g(\varDelta'_g - \hat{\varDelta}'_g)^2}{\sum_g w_g(\varDelta'_g - \bar{\varDelta}')^2}, \qquad \bar{\varDelta}' = \sum_g w_g \varDelta'_g. \tag{S7}$$

**Normalized inverse rank.** The normalized inverse rank (NIR) [25] measures whether a prediction is closer to the measured response of its own combination than to those of other combinations. We computed the Euclidean distance from $\hat{\varDelta}$ to the measured responses of all $P$ combinations of the dataset, or of the cell type in D-SPIN. The rank of the prediction is 1 plus the number of combinations whose measured response is strictly closer to $\hat{\varDelta}$ than its own, and NIR $= 1 - (\text{rank} - 1)/(P - 1)$. NIR is 1 when the own combination is the closest and 0 when it is the farthest.

**Comparison with the additive model.** To compare a method with the additive model on individual combinations, we counted for each metric the test combinations on which the method scored strictly better than the additive model, with ties counted as half. The win fraction is this count divided by the number of test combinations.

##### 1.8 Selection of sub-additive example combinations in Norman and D-SPIN

For each of Norman and D-SPIN, we selected one sub-additive example combination from the seen2 test combinations without using any model prediction. For each combination of perturbations $A$ and $B$, we computed the suppression ratio, the magnitude of the sum of the measured single-perturbation responses divided by that of the measured combination response, $\|\delta(A) + \delta(B)\|/\|\varDelta(A, B)\|$. A ratio above 1 indicates a sub-additive combination. Within one interaction class per dataset, we chose the combination with the highest suppression ratio.

For Norman, the class was suppression as annotated by Norman et al. [1], which contains 12 seen2 combinations. CEBPE + PTPN12 had the highest suppression ratio, 2.04, and the median across all 131 Norman seen2 combinations was 1.08. It also had the lowest fraction of the response explained by the additive model among the suppression combinations.

Interaction classes are not annotated for drug combinations. For D-SPIN, we defined redundant combinations as those whose two single-drug responses had a cosine similarity of at least 0.8. This held for 21 of the 188 seen2 combinations across the four cell types. Bosutinib + Dasatinib in CD8 T cells had the highest suppression ratio, 1.89, and the median across all 188 combinations was 1.31. The same combination was selected for any cosine threshold between 0.37 and 0.92. Fitting the combination response as a weighted sum of the two single-drug responses gave weights of 0.46 and 0.54, close to the average of the two responses.

For each example, the gene-level comparison used the eight genes with the largest absolute measured change in the combination.

##### 1.9 Analysis of the Wessels screen

**Prediction of the unmeasured pairs.** We trained GAIN separately on the transcriptome, with 9,391 genes, and on the 22 surface proteins (Datasets). In both models, each perturbed gene was represented by its loadings on the top 128 principal components of the log-normalized control cells, as for Norman CRISPRa (GAIN). The output subspace was computed from the training responses, with 50 dimensions for the transcriptome and 21 for the surface proteins. To assess accuracy, we randomly divided the 158 measured pairs into five folds and trained on all 28 single-gene perturbations and the pairs of the other four folds, as in the seen2 setting (Evaluation settings). We compared GAIN with the additive model and with a mean baseline, which predicts the mean response of the training pairs for every test pair, using the Pearson correlation, WMSE, $R^2_w(\Delta)$ and NIR (Evaluation metrics). For NIR, the candidates were the measured responses of all 158 pairs. To predict the 167 unmeasured pairs, we trained each model on all 186 measured perturbations.

**Synergy and dampening of CD11b.** Following Wessels et al. [8, Fig. 4c], we measured the interaction of a pair on CD11b by its deviation from the average linear relationship between the CD11b change of the pair and the mean CD11b change of its two genes knocked down alone. We fit this relationship by least squares across pairs, $y = a + bx$, where $x$ is the mean of the two single-gene changes and $y$ is the change of the pair. The deviation of a pair is $y - (a + bx)$, so that a positive deviation indicates synergy and a negative deviation indicates dampening. For the held-out pairs, we fit the line on the training pairs of each fold and computed the deviation from the measured change, from the GAIN prediction and from the additive prediction, which is the sum of the two single-gene changes. We compared each predicted deviation with the measured one by the agreement of their signs and by the Spearman correlation across pairs. We also report the mean absolute error between the predicted and measured CD11b changes of the held-out pairs. For the unmeasured pairs, we fit the line on all 158 measured pairs and computed the deviations from the GAIN predictions. We used the protein model for CD11b and the transcriptome model for ITGAM, the gene encoding CD11b.

**Classes of genetic interaction.** We followed the genetic interaction analysis of Wessels et al. [8], which was adapted from Norman et al. [1]. We used the 1,530 genes with the highest mean count per cell, the same number as in the original study, and divided the response of each gene by its standard deviation across the control cells. For a pair of genes $a$ and $b$, we fit the response to the pair as a weighted sum of the two single-gene responses, $\Delta(a,b) = c_a\delta(a) + c_b\delta(b) + \varepsilon$, by robust regression with the Huber loss. From this fit we computed six features: the distance correlation between $c_a\delta(a) + c_b\delta(b)$ and $\Delta(a,b)$, which measures the fit; the dominance $|\log_{10}(c_a/c_b)|$; the magnitude $(c_a^2 + c_b^2)^{1/2}$; the distance correlation between the two single-gene responses together and $\Delta(a,b)$; the distance correlation between $\delta(a)$ and $\delta(b)$; and the equality of contribution, the smaller of the distance correlations of $\delta(a)$ and $\delta(b)$ with $\Delta(a,b)$ divided by the larger. As in the original study, we used only pairs for which the two single-gene perturbations and the pair were each profiled in at least 25 cells, which left 142 measured pairs. We standardized each feature across these pairs and grouped them into four classes by hierarchical clustering with Ward linkage on Euclidean distances. For the unmeasured pairs whose two single-gene perturbations were each profiled in at least 25 cells, we computed the same features with the GAIN prediction in place of $\Delta(a,b)$, standardized them with the means and standard deviations of the measured pairs and assigned each pair to the class with the nearest centroid.

#### Supplementary figures

| Model | Prediction level | Perturbation representation | Norman CRISPRa<br>genetic<br>scRNA-seq | D-SPIN<br>drug<br>scRNA-seq | ComboSciPlex<br>drug<br>scRNA-seq |
|---|---|---|---|---|---|
| GAIN | Population | Control-cell PCA gene loadings; Morgan fingerprints | ● | ● | ● |
| Additive | | Measured single-perturbation responses | ● | ● | ● |
| GEARS | Single cell | Gene Ontology gene graph | ● | — | — |
| Scouter | | GenePT gene text embeddings | ● | — | — |
| Geneformer | | In silico gene up-regulation in rank tokens | ● | — | — |
| CellFlow | | ESM2 protein embeddings; Morgan fingerprints | ● | ● | ● |
| ChemCPA | | Molecular (SMILES) drug embedding | — | ● | ● |

● evaluated in this study — not applicable

**Supplementary Fig. 1.** Models and datasets evaluated in this study. The table lists the prediction level and perturbation representation of each model, and the datasets on which each model was evaluated. Single-cell models were applied only to the perturbation type they support.

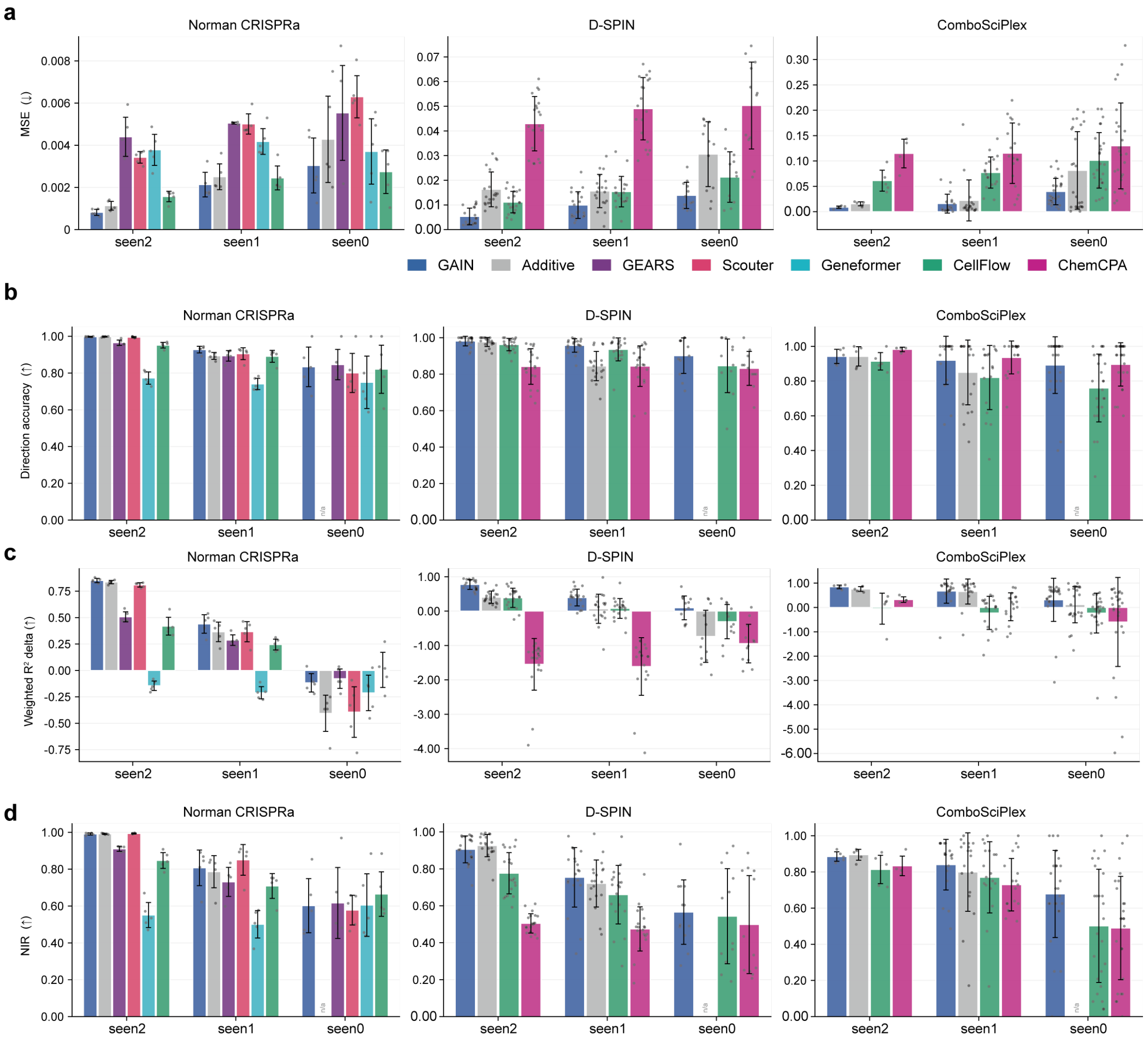


**Supplementary Fig. 2.** Additional benchmark metrics. Bar plots show the MSE (**a**), direction accuracy (**b**), weighted $R^2_\Delta$ (**c**) and NIR (**d**) of each model on Norman, D-SPIN and ComboSciPlex in seen2, seen1 and seen0, in which both, one or neither of the two single-perturbation responses are in the training set. The direction accuracy is the fraction of the 20 genes with the largest measured change whose predicted change has the same sign. The weighted $R^2_\Delta$ measures how much of the response specific to a combination is predicted, with genes weighted by how specifically they respond to the combination. NIR (normalized inverse rank) measures how close a prediction is to the measured response of its own combination relative to those of the other combinations, and ranges from 0 for the farthest to 1 for the closest. Arrows indicate favorable directions. Bars show the mean and error bars the standard deviation across folds, and dots show individual folds. For D-SPIN, each cell type and fold is counted separately. The additive prediction is zero in seen0, so its direction accuracy and NIR are marked n/a.

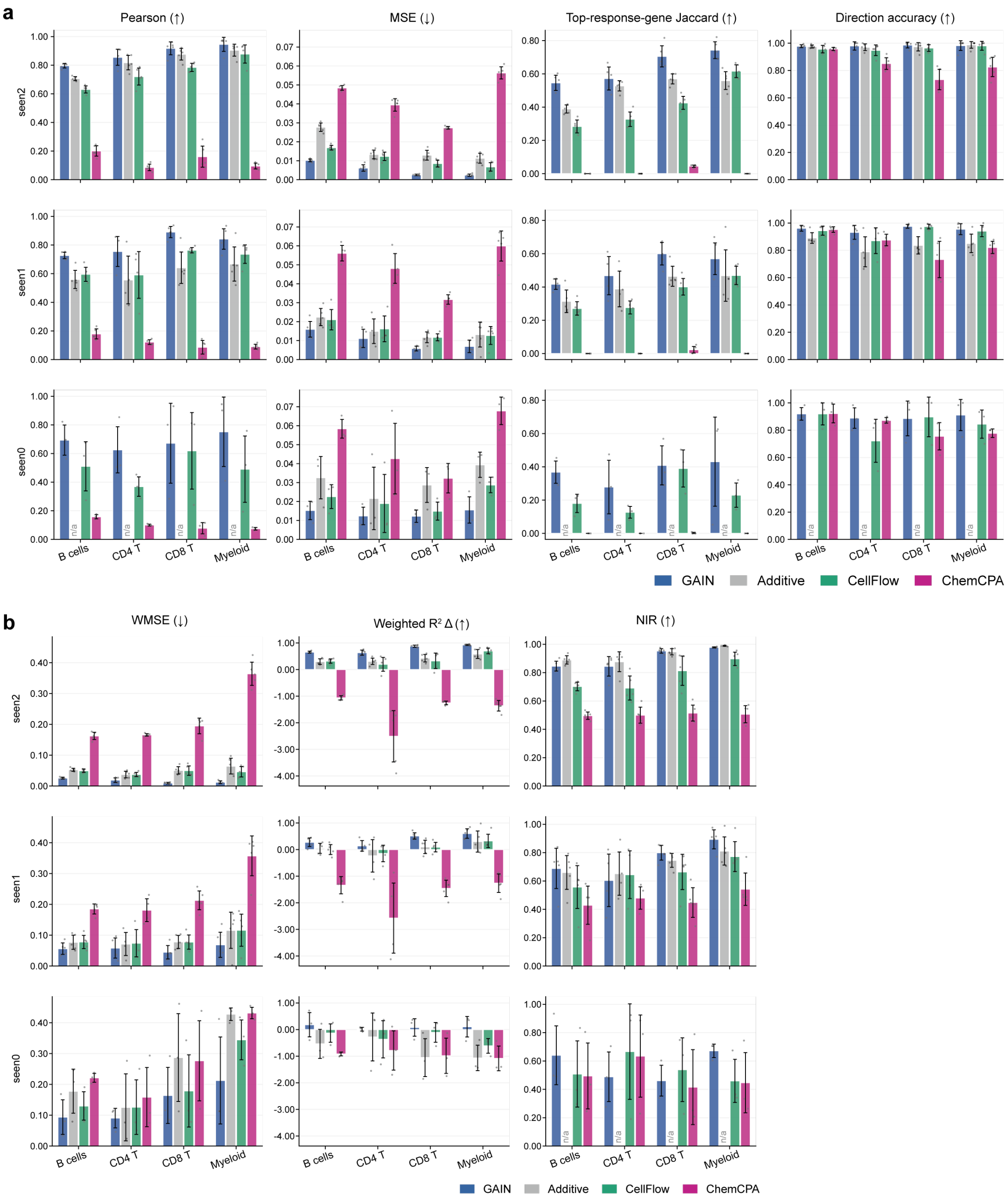


**Supplementary Fig. 3.** Benchmark on each D-SPIN cell type. **a**, Bar plots show the Pearson correlation, MSE, top-response-gene overlap and direction accuracy of each model on B cells, CD4 T cells, CD8 T cells and myeloid cells in seen2, seen1 and seen0. **b**, Bar plots show the WMSE, weighted $R^2_\Delta$ and NIR for the same cell types and settings. Arrows indicate favorable directions. Bars show the mean and error bars the standard deviation across folds, and dots show individual folds.

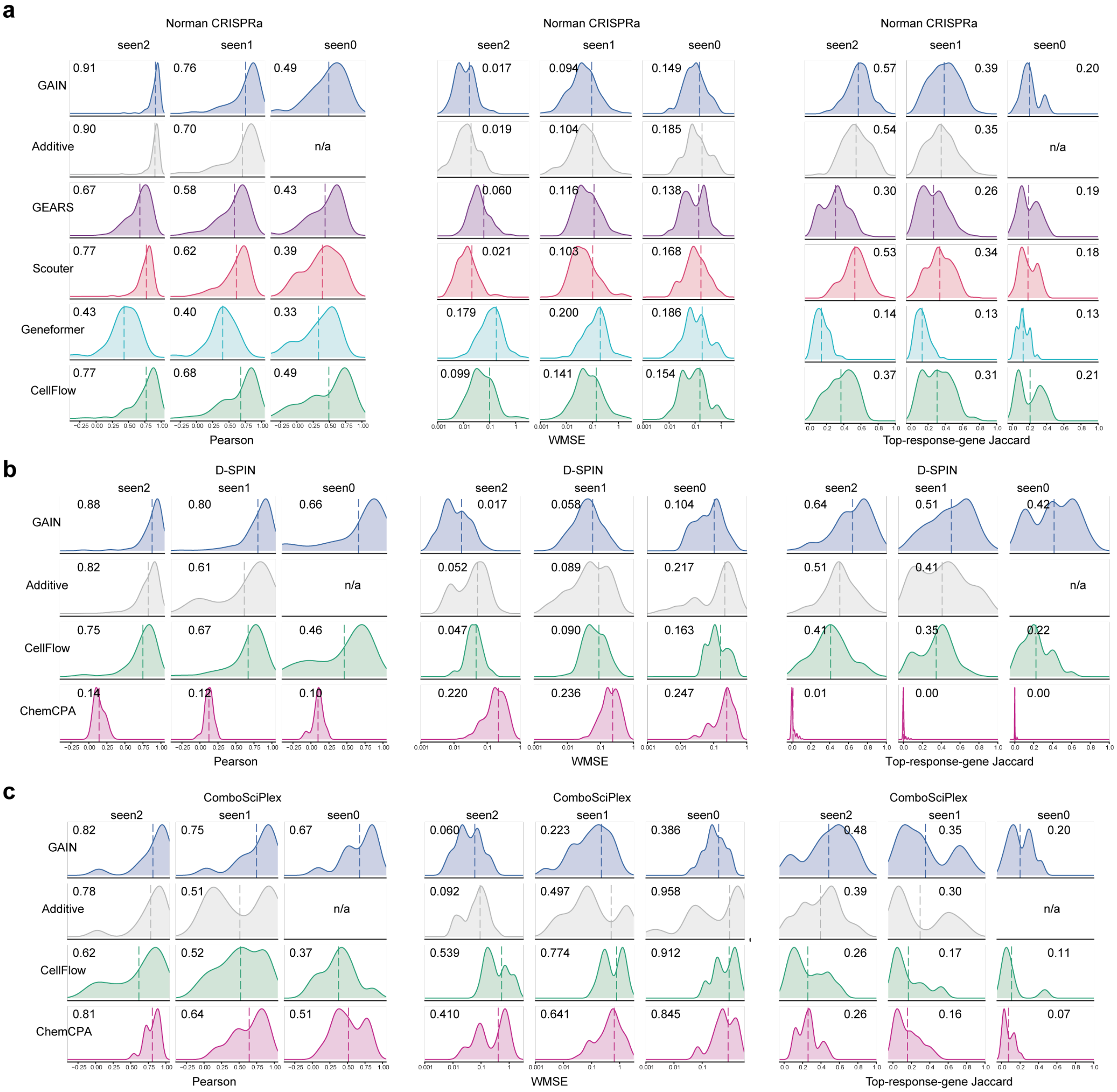


**Supplementary Fig. 4.** Distributions of per-combination scores. Density plots show the distributions of the Pearson correlation, WMSE and top-response-gene overlap over test combinations, pooled across folds, for each model in seen2, seen1 and seen0 on Norman (**a**), D-SPIN (**b**) and ComboSciPlex (**c**). Dashed lines and numbers represent the mean. For D-SPIN, each cell type is counted separately. The additive prediction is zero in seen0, so its Pearson correlation and top-response-gene overlap are marked n/a.

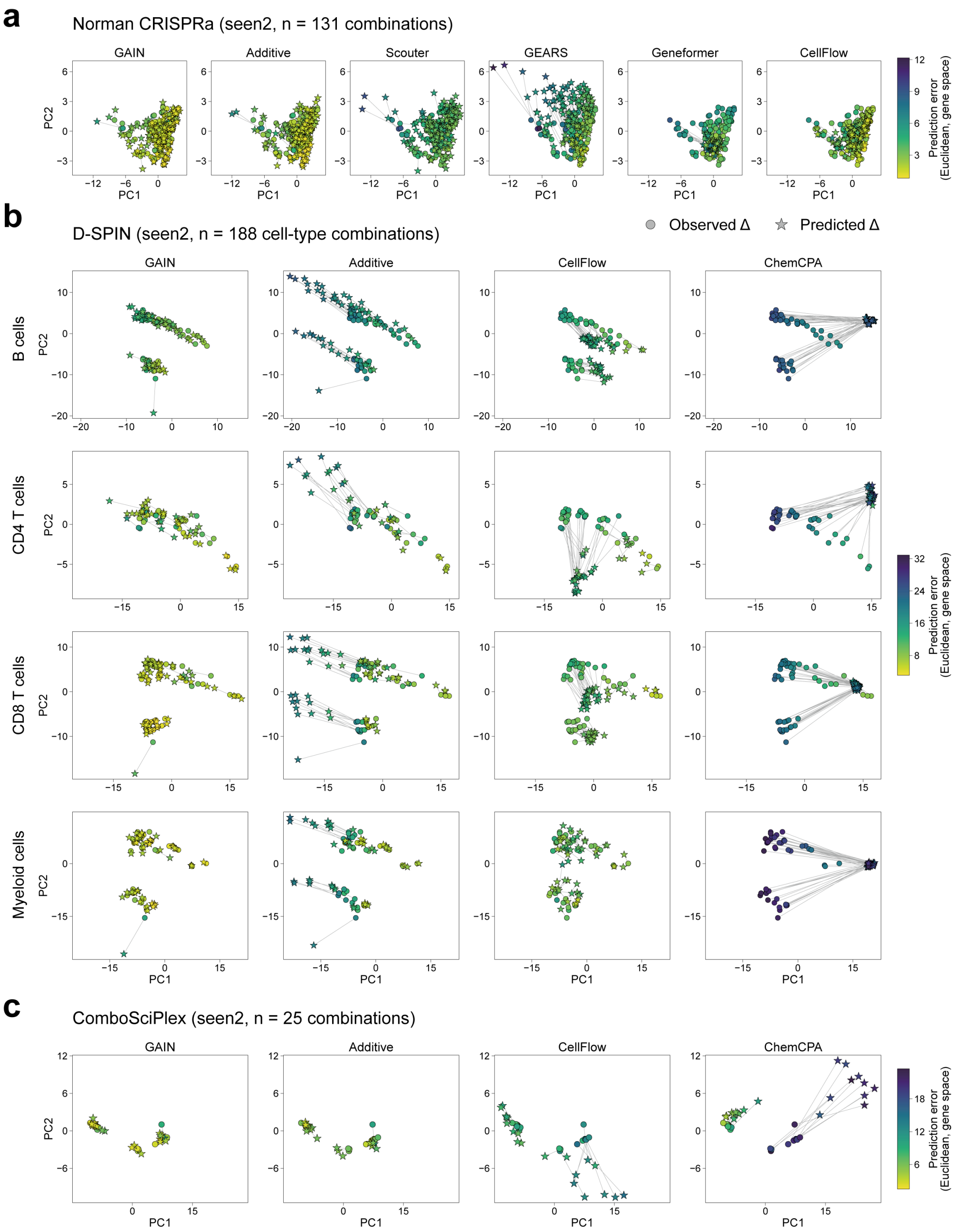


**Supplementary Fig. 5.** Measured and predicted responses in the space of the two leading principal components. Scatter plots show the measured and predicted responses of the seen2 test combinations on their first two principal components for Norman (**a**), D-SPIN (**b**) and ComboSciPlex (**c**). The components were computed jointly from the measured responses and the predictions of all models, separately for each D-SPIN cell type. Circles show measured and stars predicted responses, and gray lines connect the two for each combination. Color shows the Euclidean distance between the predicted and measured responses in gene space.

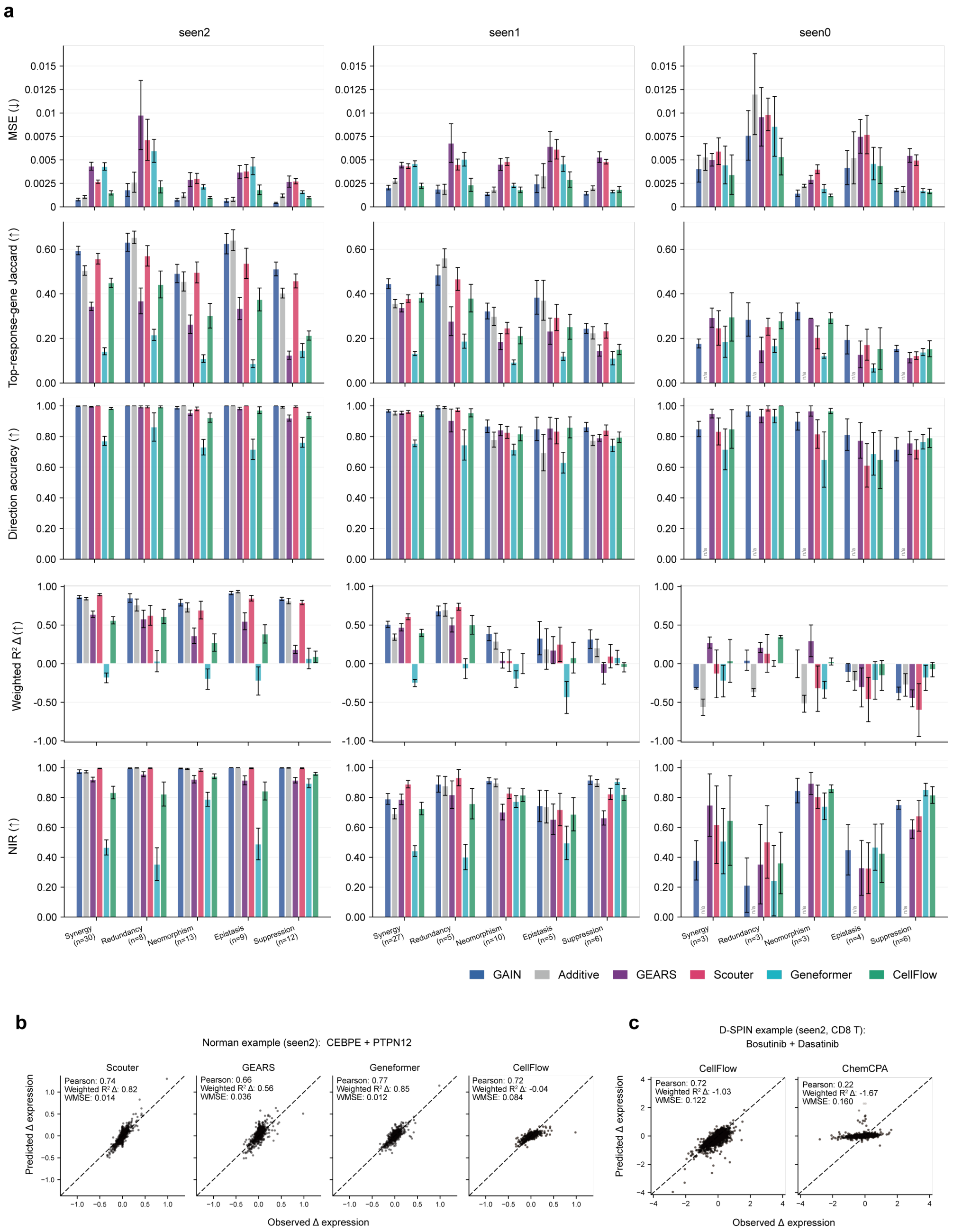


**Supplementary Fig. 6.** Performance by interaction class and the other models for the two examples. **a**, Bar plots show the MSE, top-response-gene overlap, direction accuracy, weighted $R^2_\Delta$ and NIR of each model on Norman combinations grouped by interaction class in seen2, seen1 and seen0. Arrows indicate favorable directions. Bars show the mean and error bars the standard error of the mean across combinations, and $n$ is the number of combinations in each class. **b**, Scatter plots show the predicted against the observed responses of all genes to CEBPE + PTPN12 in Norman (seen2) for Scouter, GEARS, Geneformer and CellFlow. **c**, Scatter plots show the predicted against the observed responses of all genes to Bosutinib + Dasatinib in D-SPIN CD8 T cells (seen2) for CellFlow and ChemCPA. In **b** and **c**, each dot is one gene, and values give the Pearson correlation, weighted $R^2_\Delta$ and WMSE of each prediction.

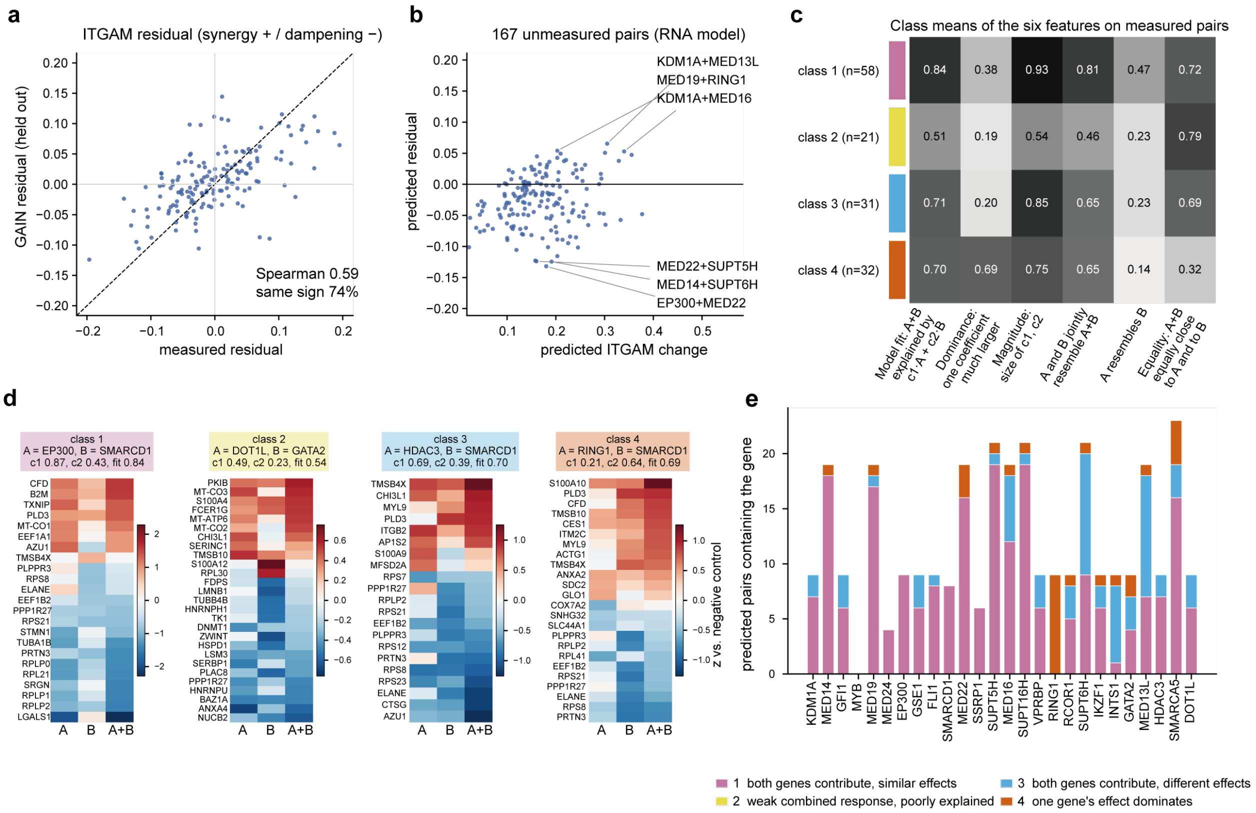

**Supplementary Fig. 7.** ITGAM residuals and interaction classes in the leukemia differentiation screen. **a**, The scatter plot shows the ITGAM residual predicted by the GAIN RNA model against the measured residual for each held-out pair. The residual of a pair is its deviation from the average linear relationship between the ITGAM change of a pair and the mean ITGAM change of its two single-gene knockdowns. Values give the Spearman correlation and the percentage of pairs whose predicted and measured residuals have the same sign. **b**, The scatter plot shows the predicted residual against the predicted ITGAM change for the 167 unmeasured pairs. Pairs with the largest positive and negative residuals are labeled. **c**, The heatmap shows the mean of the six regression features in each interaction class over the measured pairs, and $n$ is the number of measured pairs in each class. **d**, Heatmaps show the responses to the knockdown of gene A, gene B and both for one measured pair per class, namely the pair closest to the class centroid, over the union of the 10 genes with the strongest response to each perturbation. Values are z-scores relative to negative control cells. **e**, The bar plot shows the number of unmeasured pairs containing each gene, colored by predicted class.
